# A next generation space antimicrobial: assessing microbial activity and reduction across the International Space Station

**DOI:** 10.64898/2026.01.10.698798

**Authors:** Jason W Armstrong, Olivia AK Jessop, David Corporal, Iain S Koolhof, Sung-Po R Chen, Nicholle R Wallwork, Reuben Strydom, Mark E Wilson, Darren S Dunlap, Ariel C Muldoon, Michael Monteiro

**Affiliations:** The Boeing Company, Washington, United States of America; Australian Institute for Bioengineering and Nanotechnology, The University of Queensland, Brisbane, Queensland, Australia

## Abstract

Microbial contamination in spacecraft poses a threat to crew health and operational integrity. Currently, microbial reduction aboard the International Space Station (ISS) relies on rigorous cleaning, which is time-consuming for astronauts. This study analyzes the ISS microbiome and applies a novel quaternized antimicrobial polymer coating to assess its effectiveness in mitigating microbial contamination and its durability over prolonged periods of interaction. The polymer was coated onto five material types representing common objects interacted with by the crew. Each material had six coupons placed on six placards in various ISS locations, while three placards remained on Earth to accumulate microbes from passive and direct transfer. The trial lasted six months, utilizing microbiological cell cultures and genomic analysis to assess bacterial communities and identify species. Among the six placards on the ISS, only one had sufficient viable bacteria to assess the polymers efficacy. On this placard, the polymer coating was shown to decrease culturable bacterial surface loads by 1.39 log_10_ (equal to a 95.9% reduction) compared with uncoated surfaces. Coated surfaces exhibited less genetic material and lower microbial species richness. Additionally, the polymer coating proved durable, persisting on ISS surfaces throughout the six-month period. This study demonstrates that the antimicrobial polymer coating can actively reduce microbial loads and remains on surfaces in zero gravity environments. Its application is likely beneficial when applied alongside other mitigation activities for viral and bacterial contamination, with potential uses in long-duration spaceflight to enhance crew health and maintain spacecraft integrity. We also generated data showing the lack of microbial presence on the ISS.

## Introduction

The internal closed system of spaceflight environments provides a series of challenges in regards to microbial infection and microbe contamination (bacteria and fungi in particular) which can compromise materials, life-support systems and human health in what is an immunocompromising environment[1,2]. Microbiological risk factors for astronaut health include: food, drinking water, air, surfaces, payloads, animals, crew members, and personnel in close contact with the astronauts[3]. Potentially harmful microbial growth could also be exacerbated by mutation rate, replication and biomass of microbes in microgravity[4]. Although the effect of spaceflight on surface microbes has been minimally studied, it has been proposed that spaceflight does not dramatically alter the growth of surface microbes compared to Earth[5] with the caveats it does alter resistance to alkaline cleaning products[6] given environmentally sampled bacterial genomes from the International Space Station (ISS) showed increased DNA repair activity, enhanced metabolism through mobile genetic elements, and furthermore evolution of potential pathogenic capabilities. Novel species have been identified in space[7], supporting the idea that the niche environment does promote growth of bacteria differently than on Earth[8–10].

The importance and severity of notable spaceflight events relating to microbial growth in flights have been well summarized and include damage to equipment function^11^, and several instances where uncontrolled microbial growth has threatened crew safety. For example, the Internal Active Thermal Control System (IATCS) on the ISS experienced uncontrolled planktonic bacterial and biofilm growth in the early years of the program[11]. This led to biofouling of various components and subsequent decreased flow rates, reduced heat transfer, initiation and acceleration of corrosion, and enhanced mineral scale formation.

Microbial spread can occur by three primary modes: aerosol, close contact, and fomite transmission by surface or object direct contact[12]. The risk of aerosolized particles can be somewhat mitigated by the engineering of air filtration and circulation systems[10]. Preventing fomite transmission, however, currently relies on extensive cleaning at the expense of increased time in labor, extra payload mass/volume, cost, and indirect associated hazards[4]. Currently, cleaning aboard the ISS is performed weekly at a minimum[13]. Further, the cleaning methods vary in their efficacy, application effectiveness, pathogen deactivation time, and longevity, and may not be sufficient to inactivate or remove potential pathogenic or environmentally harmful organisms. In comparison, a persistent, fast acting antimicrobial that can be applied to surfaces would be useful in closed environments in space and as a public health measure on Earth in mitigating disease risks in a variety of settings (e.g., food industry, public transportation, etc).

Antimicrobial polymers applied to surfaces provide a useful means to control microbial growth including those of potentially pathogenic origin[14]. They have potential utility in preventing the formation of biofilms, relevant to hardware surfaces and regulated water systems aboard the ISS[13,15]. Previous evaluation of polymers to the ISS has assessed their hydrophobicity mode of action, and physical application method[16]. Here we discuss the antimicrobial impact of our novel quaternized, coumarin-functionalized antimicrobial polymer coating which provides a nanomechanical membrane rupturing mode of action (resulting in cell lysis), embedded microbe specificity and uses as a persistent antimicrobial application in both spaceflight and terrestrial settings[14,17]. The coating consists of a polymer that has broad anti-bacterial and anti-viral activity, and most notably it allows for specific functionality to be built in to target multiple microbial strains at once[17]. The nanoparticles respond to touch, moisture, temperature and pH to achieve microbial lysis. Further, it is fast acting, durable over a long-term period, water-resistant and has a degree of self-cleaning. While ground studies have shown the efficacy of this technology, testing this polymer on the ISS across two missions was aimed at understanding its function in the varied parameters of spaceflight including: microgravity, higher radiation levels, close quarter living and the material types used in spacecraft cabins. Building from an initial flight test in 2021[18], this study aims to assess the effectiveness of this antimicrobial polymer on microbes found in the ISS through bacterial quantification and identification via culture and complementary genetic methods, as well as durability on representative high touch surfaces[19].

## Materials and Methods

The antimicrobial coating used here was previously flown to ISS on SpaceX-21 in December 2020 and returned to the ground on SpaceX-22 on July 2021[18]. The efficacy of the polymer coating was assessed for bacteriological and fungal growth. Microbial presence was less than expected which was at first thought likely due to the touching method not including a re-inoculation procedure and the delay in sample post-flight analysis given logistical challenges caused by the COVID-19 pandemic. While on this mission, post-landing pandemic staff limitations and site access impacted post-flight analysis, so data interpretations were limited but it did suggest relatively low contamination levels for ISS, at least in the assessed area with touch interactions as a variable.

The aim of the second flight, flown from November 2023 to April 2024, was to test the efficacy of the polymer coating against microbes in various locations on the spacecraft, gauge general microbial contamination levels and assess the durability of the polymer in prolonged spaceflight.

### Experimental Design

Nine placards were prepared in total, six were placed in four locations on the ISS: three placards in the galley (Node 1), one in the exercise area (Node 3), one in the hygiene area (Permanent Multi-Purpose Module), and one in the toilet (Node 3); and three placards were left on ground (Figure 1 & Table 1). These locations were chosen as the most likely areas to experience a larger presence of microbes when compared to other areas on the station[4]. Each placard carried five material types, with six coupons per material type: seat belt strap, cargo transfer bag (CTB) fabric, medical gauze, tray table material, and E-leather. On each placard, three coupons per material were coated with the antimicrobial polymer and an adjacent three coupons were left uncoated as a matched control, where coated/uncoated pairs were fixed next to each other (Figure 2). Thus, each placard provided matched coated/uncoated comparisons for the five materials.

**Figure 1.**
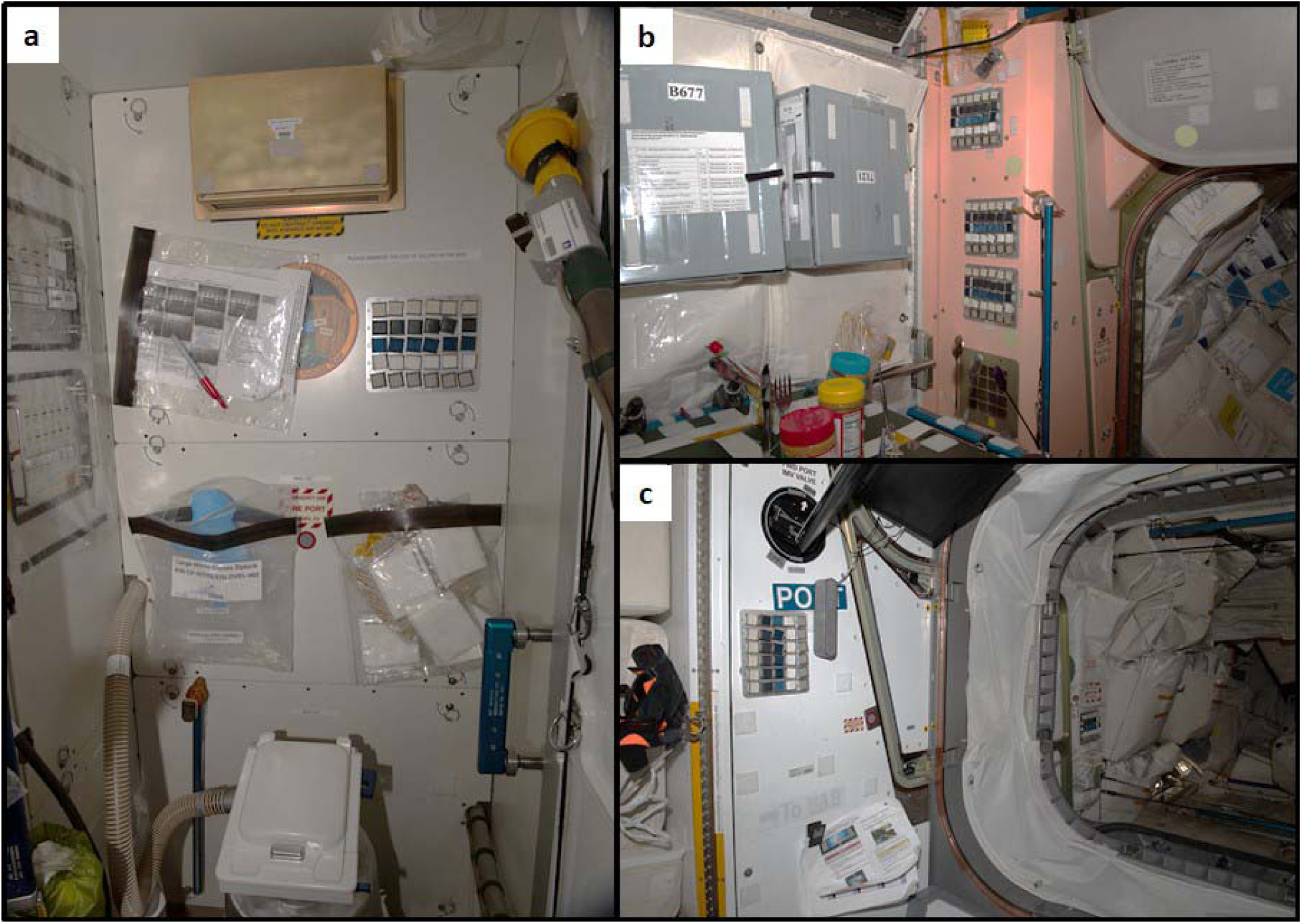
Placard locations on the ISS: a, Toilet (Node 3); b Galley (top – Galley 3, middle – Galley 2, Bottom – Galley 1) (Node 1); and c Exercise area (left) (Node 3) & Hygiene area (right) (Permanent Multipurpose Module).

**Table 1.**
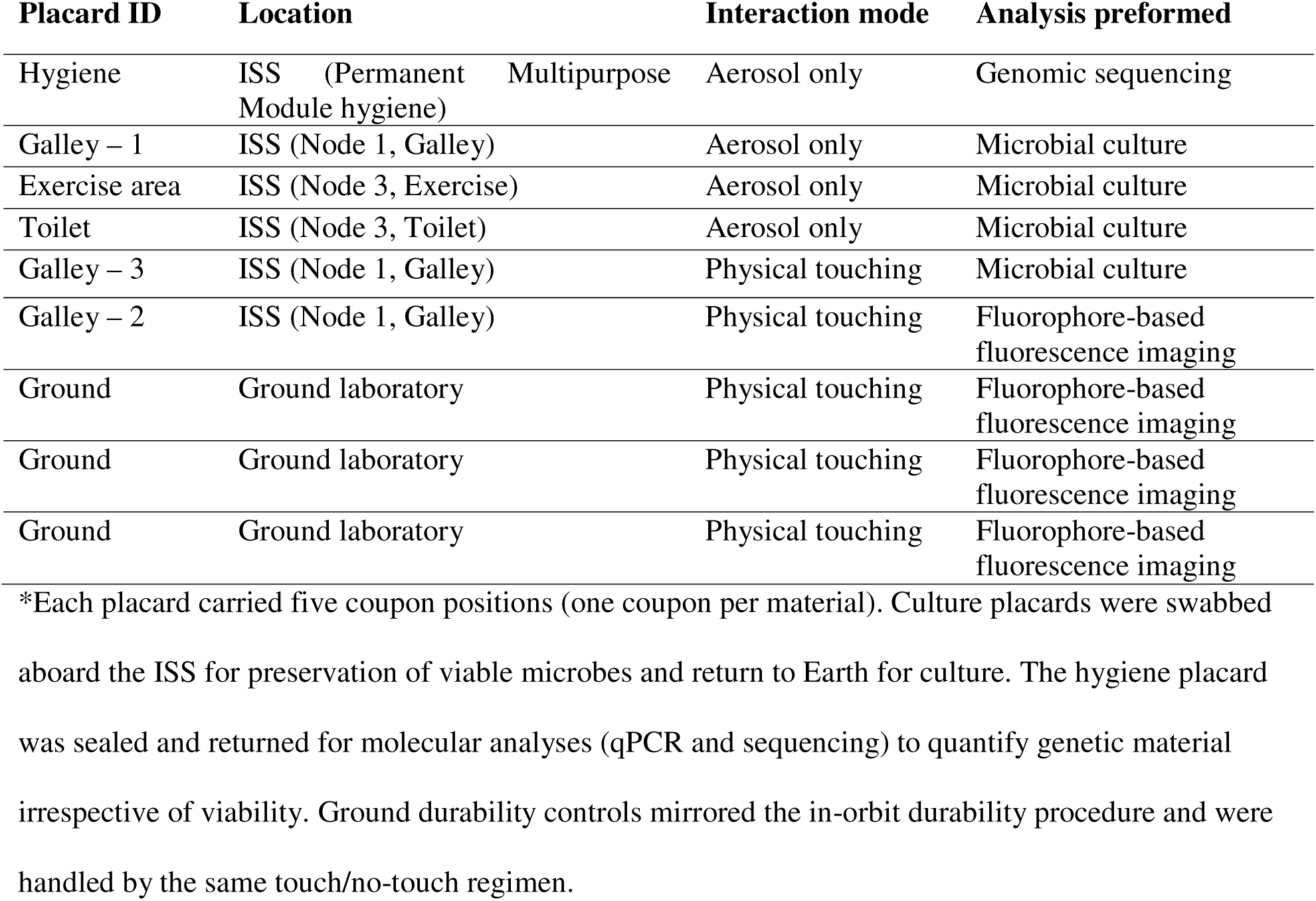
Placards distributed on the ISS and ground for microbial load and durability assessment.

**Figure 2.**
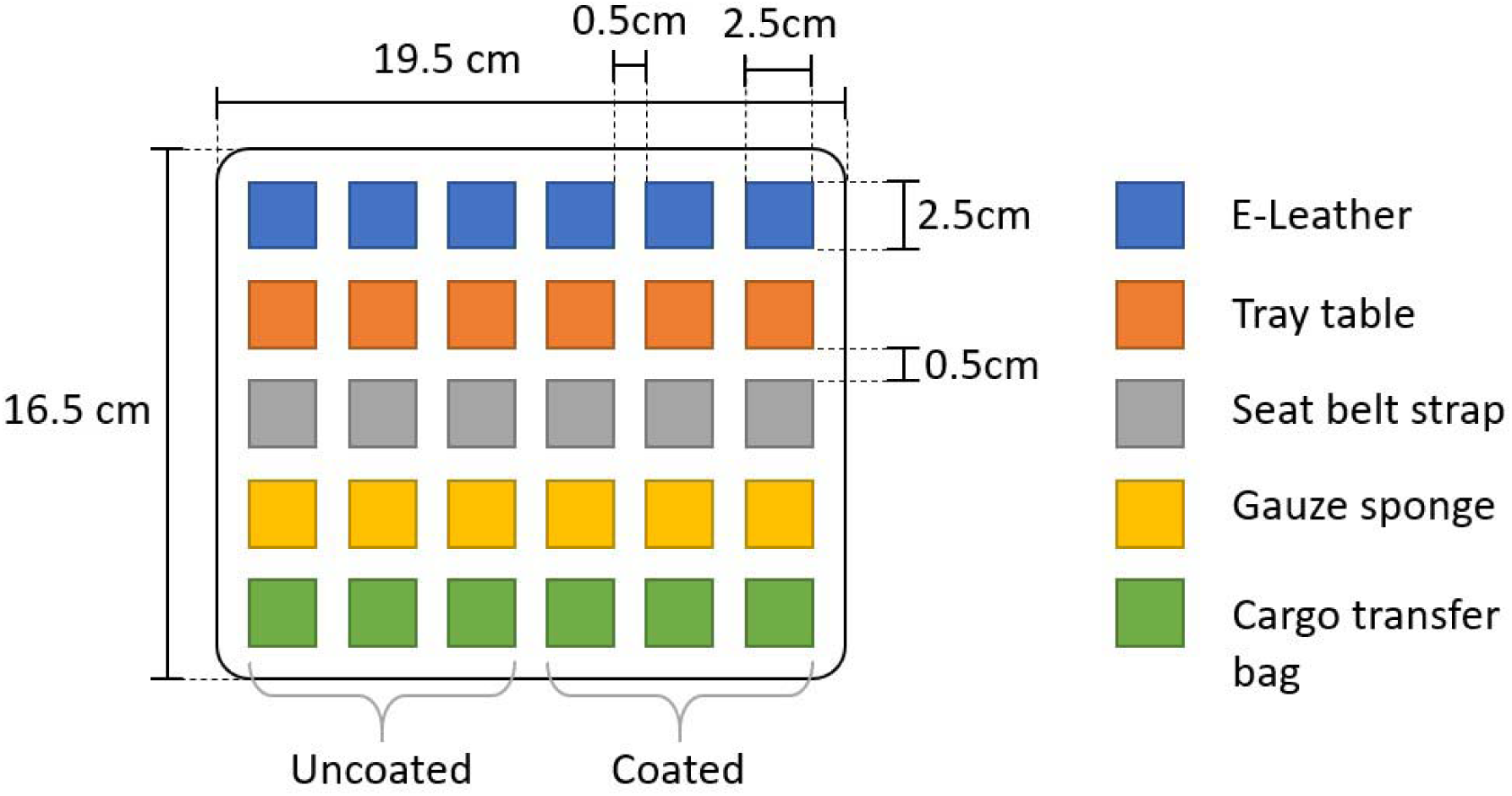
Coupon layout among individual placards aboard ISS.

Each coupon on a placard is treated as an independent replicate. In total 270 coupon positions were prepared across all locations (6 ISS placards × 5 material types × 6 coupons = 180; 3 ground placards × 5 material types × 6 coupons = 90). Owing to logistical constraints for two of the placards undergoing microbial culturing (Gallay – 3 and the Toilet placard) only four out of the six coupons per material type were sampled for analysis (two coated and two uncoated with the polymer). Culture-based analyses were possible for only the Galley – 3 placard; microbial culture statistics therefore used only the Galley-3 coupon set (*n* = 20). Genomic analyses used the hygiene placard coupons (*n* = 30) and durability imaging used the designated durability placards and ground controls (*n* = 120).

On each placard, polymer coated and uncoated coupons were paired and placed adjacent to one another to provide matched (paired) biological comparisons for each material at that location. Technical splits were performed during the microbial analysis where the culture plating included duplicate platings for each dilution from a given swab, and qPCR was run in triplicates across three dilutions (neat, 1:5, 1:10) per DNA extract (detailed process described below). For downstream statistical analyses, microbial culture colony forming units (CFU) per cm^2^ were averaged duplicate plate counts to yield a single count per coupon; for qPCR we used the mean Ct of technical replicate wells (setting undetermined wells as missing) and converted to gene copies using the standard curve. Samples with no technical replicate producing Ct were recorded as non-detected, and technical replicates were not treated as independent observations.

To evaluate durability, fomite transfer, and both culture-based and molecular microbial detection, specific placards were designated for each analytical approach (Table 1). Culture-based microbial enumeration and isolate identification were performed on the microbial culture placards (fomite and selected aerosol placards). Genomic sequencing, quantitation, and community profiling (qPCR and 16S amplicon sequencing) were performed on the hygiene placard. Polymer durability was assessed on designated durability placards using fluorophore-based fluorescence imaging (see Table 1).

The hygiene placard was deployed as an aerosol (no-touch) sampling unit to characterize airborne/environmental deposition of microbial genetic material independent of direct crew contact. This was to test whether the molecular (qPCR and 16S amplicon sequencing) assays would detect low-biomass aerosol deposition with greater sensitively than microbial culturing; therefore, the hygiene placard was returned sealed for DNA-based analyses. This placard was intended to permit direct comparison between aerosol-deposited communities and those from fomite (touch) placards, not to serve as a negative control.

#### Experimental Hardware and Sample Preparation

The antimicrobial polymer coating used in this study was a quaternized, coumarin-functionalized polymer dispersion synthesized by post-polymerization modification to introduce antimicrobial cationic groups and a fluorophore for durability imaging (see Supporting Information 1). Coupons were prepared (2.5 × 2.5 cm) and sterilized using an appropriate method per material: autoclave (121 °C) for autoclave-tolerant items, UV-C exposure within a laminar flow cabinet for sensitive items, or 70% ethanol spray where compatible. The antimicrobial polymer dispersion was sprayed onto designated coupons at ∼7 cm distance and air-dried; the spray–dry cycle was repeated four times to achieve the intended coating. Coated coupons were paired adjacent to uncoated control coupons on placards using Velcro. Negative controls (three packaged placards sealed prior to launch and blank extraction controls) were processed in parallel through culture and molecular analysis; sequencing and qPCR results were screened against these controls and any taxa/features present in negative controls were flagged and considered in downstream interpretation. Dedicated sequencing blanks were included, including during DNA extraction, PCR amplification, and library preparation. These controls were processed in parallel with the ISS samples and were used to monitor for and assess background contamination introduced during sample handling and molecular processing. Features detected in blank controls were considered during downstream interpretation of low-biomass samples.

All hardware assembly was performed under aseptic conditions. On the ground, matched durability placards were split into control and treatment groups: the treatment group was touched three times per week over 24 weeks according to the ground touch protocol; controls were left undisturbed for baseline fluorescence comparisons. An in-depth description of the sterilization methods and the preparation of the polymer mixture prior to coating can be found in the methods section of the supporting information 1.

#### Placard placement, Crew Interaction, and Sampling

Placards were positioned at representative locations aboard the ISS expected to receive fomite or aerosol deposition as summarized in Table 1. For fomite (touch) placards, crew performed a systematic touch regimen approximately every 48 hours (subject to crew time availability) to deposit microbes via contact; both coated and uncoated coupons on these placards were touched. For durability placards, a subset of coupons was touched repeatedly while others were left untouched to assess polymer retention under use. Aerosol (no-touch) placards were left unhandled to accumulate environmental deposition. Placards intended for culture were swabbed across their full surface on orbit using Amies agar gel transport swabs to preserve viable microbes during return (kept at 4°C during return). Crew swabbing technique was verified with project personnel. The hygiene placard was sealed and returned without culture swabbing; the coupons were processed on return for DNA extraction and molecular assays.

### Microbial Culture Enumeration

Swabs were vortexed in 3 mL phosphate buffered saline. Ten-fold dilutions were performed on each sample from 10^-1^ to 10^-2^ in sterile phosphate buffered saline. Microbial enumerations were performed by plating 100 µL of these dilutions on 2 different media including a non-selective Tryptic Soy Agar (TSA) media for heterotrophic bacteria and a selective media for fungi (Sabouraud dextrose agar (SDA) with 0.025 g/L or 0.005% w/v chloramphenicol) per Standard Methods for the Examination of Water and Wastewater[20]. Media was selected to match current methods for monitoring microbial contamination on cabin surfaces on ISS[21].

Each sample dilution was performed in duplicate. TSA plates were incubated for heterotrophic bacteria for 7 days at 37°C and colonies were counted at 2 days. Fungi plates were incubated at 25°C room temperature for 5 days and counted colonies at 2 days and 5 days.

#### Microbial Isolate Identification

Identifications were performed by isolating each different colony morphology on a plate in pure culture. Bacterial identifications were performed to genus and species using fatty acid methyl ester gas chromatography and 16S sanger sequencing. Fungal isolates were identified using macroscopic and microscopic morphology. Pure bacterial cultures were identified firstly using fatty acid methyl ester (FAME) fingerprinting via the Microbial Identification System (MIDI, Inc., Newark, NJ). DNA was then extracted from microbial isolates using the Zymo Quick-DNA Fungal/Bacterial Mini-Prep Kit and amplified by using 27F/1492R full 16S universal primers for bacteria shown in S1 Table 1. Sanger sequences were trimmed for quality and assembled using BioEdit version 7.0.0. The bacterial consensus sequences were aligned against the SILVA SSU rRNA database using ARB software (http://www.arb-home.de/)[22] with a 95% minimum identify for genus-level and a 99% minimum identity for species-level identification. Bacterial isolates were also compared against the NCBI BLAST database using the same cutoff parameters. Organisms that were unable to be identified to species level but were identified to genus level were listed as “sp.” Organisms that were unable to be identified to genus level were listed as “Low match” or unidentified.

### Molecular Quantitation and DNA Sequencing

The hygiene placard was specifically designated to capture aerosol deposition (no touch) and was processed for molecular assays on return to maximize detection sensitivity for low biomass airborne deposition. Each coupon on the hygiene placard was swabbed with a flocked swab (4N6FLOQSwabs™, Thermo Fisher) pre-wetted with 5 µL nuclease-free water and sampled across the entire coupon using a standardized pattern (top→bottom, left→right, and two diagonals). Swab tips were snapped into 2 mL screw-cap tubes containing 800 µL Solution CD1 and 300 mg 0.1 mm glass beads, then lysed by bead beating for 5 min at 2000 rpm (Powerlyser 24). Tubes were heated to 65 °C for 10 min with shaking and centrifuged; lysates were processed using the Qiagen DNeasy PowerSoil Pro kit with a final elution volume of 50 µL (detailed stepwise protocol and reagent catalog numbers are provided in Supporting Information 2).

Quantitative PCR (qPCR) targeted the bacterial/archaeal 16S rRNA gene (primers 1406F/1525R; see Supporting Information 2 for sequences). Reactions (11 µL total) used 6 µL 2× QuantiNova SYBR Green Master Mix (Qiagen), 4 µL template DNA and 1 µL primer mix (0.4 µM). Each extract was assayed in technical triplicate at three dilutions (‘neat’, ‘1:5’, ‘1:10’). Cycling conditions on the ViiA 7 platform were: 95°C 2min; 40 cycles of [95°C 5s, 60°C 20s]; followed by a melt curve. Cycle threshold (Ct) values were recorded and used to estimate 16S gene copy abundance after accounting for dilution and standard-curve metrics (see Supporting Information 2 for details).

For community profiling, we amplified the V3–V4 region of the 16S rRNA gene using Illumina-adaptered primers (341F/806R with overhang adapters). PCR used NEBNext Ultra II Q5 Mastermix to generate ∼465 bp amplicons; amplicons were purified with Agencourt AMPure XP beads at 0.8× ratio, indexed using the Illumina Nextera XT indexing kit, pooled equimolarly and sequenced on an Illumina MiSeq (V3 2×300 bp). Per-sample raw read minimum to pass QC was 10,000 reads. Read processing removed primer sequences with ‘cutadapt’ (v2.4), trimmed low quality bases with ‘Trimmomatic’ (v0.39; SLIDINGWINDOW:4:15, CROP:250, MINLEN:250), and denoised with DADA2 implemented in QIIME2 (ver. 2019.10). Only forward reads were used for amplicon analysis because reverse reads failed to meet the minimum overlap/quality thresholds across samples for reliable pairing; the denoising and chimera removal employed DADA2 in QIIME2. Taxonomy was assigned by BLASTing features against the SILVA SSU database (release 138, 99% clustering). Computational pipeline parameters and downstream filtering thresholds are provided in Supporting Information 2.

### Polymer Durability Assessment

The polymer was formulated with an adjuvanted fluorophore to allow nonDdestructive detection of remaining coating [17]. FluorophoreDtagged coupons were imaged under UVDA (365 nm) excitation using a standardized camera setup at biweekly intervals for ground controls and preD/postDflight for ISS durability placards where feasible. Image intensity (mean gray value) within defined coupon regions of interest was quantified using ImageJ. Prior ground validation confirmed negligible quenching of the fluorophore under tested lighting regimes. Durability comparisons were made between touched and untouched coupons, and between ISS and ground controls.

### Statistical Analysis

Summary statistics were used to assess qualitative differences in log_10_ microbial CFU/cm^2^ between material type and the application of the polymer, displaying count information, the mean (x□) and standard deviation (σ) between each group. An analysis of variance (ANOVA) was used to determine whether there was a difference in mean CFU/cm^2^ between the materials and polymer coating. Owing to the low number of samples per material type, there was insufficient data to conduct a statistical test on the interaction between material type and the polymer coating, thus separate univariate analysis were used.

A linear regression model of the log_10_ bacterial CFU/cm^2^ was used to examine the response from the use of the polymer coating. A constant (one log_10_) was added to bacterial CFU/cm^2^ to allow for counts to adjust for coupon that had no microbial presence. Model assumptions were checked using residual plots and assumptions of variance heterogeneity and normality of errors were reasonably met.

Differences in the microbial species communities between material type and polymer coating were conducted on the hygiene placard. Summary statistics were used to identify and quantify the relative abundance of genetic material by each taxon detected from genomic analysis described above. The relative abundance of each taxon was then visualised using Bray–Curtis dissimilarities computed on the Hellinger-transformation and used as input to Non-metric multi-dimensional scaling (NMDS) analysis [23] to characterize species communities among the samples. An analysis of similarities (ANOSIM), a non-parametric statistical test, was conducted to construct dissimilarity matrices to test whether there were statistically significant differences between the composition of microbial species from coupon material types and whether the coupon was coated or uncoated with the polymer. Multi-level pattern analysis (MPA) was then used to determine if the amount of genetic material present on the coupons significantly differed between material type and between coated and uncoated coupons. Both the ANOSIM and MPA used 9,999 permutations to assess the significance of species and group associations where relevant. Because ANOSIM may be sensitive to unequal within-group variance, we further tested homogeneity of multivariate dispersions using the polymer coating and material type as grouped centroids and evaluated significance using Permutational Multivariate Analysis of Variance (PERMANOVA, with 9,999 permutations).

Lastly, the amount of genetic material per taxon was examined using a linear model with a natural log of abundance as the response variable and the polymer coating, material type, and their interaction as explanatory variables. Model assumptions were checked using residual plots and assumptions of variance heterogeneity and normality of errors were reasonably met. However, an extreme amount of DNA for a single taxon was found on a single coupon on the E-leather material leading to the observation being treated as an outlier in the right-hand tail of the distribution of residuals. Due to the log transformation, inference is to multiplicative differences in medians. A negative binomial generalized linear model with a log link was fitted to the number of species data, with the polymer coating, material type, and their interaction included as explanatory variables, excluding the single outlier observation mentioned above. Model fit was checked using simulated residuals plots and no problems were noted.

Statistical analysis was undertaken in R (Version 4.0.3, www.r.project.org)[24], using the latest compatible versions of packages ‘MASS’[25], ‘vegan’[26], ‘ggplot2’[27], DHARMa[28], & ‘indicspecies’[29] in RStudio (Version 2023.12.1)[30].

## Results

### Microbial Growth and Identification

#### Polymer Efficacy

Of the four placards place on the ISS to assess the polymers efficacy on microbial activity, the Galley – 3 placard was the only placard to have sufficient number of viable bacteria to perform statistical analysis and is the only placard used in assessing the polymers antimicrobial efficacy. Of the remaining placards, only seven of the 70 coupons had viable bacteria (two coated and five uncoated). From the Galley – 3 placard 11 coupons had viable bacteria of which 19 of the 40 swab samples (including duplicate replicates) had detectable viable bacteria (S1 Table 2). Of the 11 coupons that had growth, only three of the coupons were from those coated with the polymer (Table 2) and of the 19 swab samples that had detectable viable bacteria, five were from polymer coated coupons and 14 from uncoated coupons.

**Table 2.** The number coupons that had detectable viable bacteria by material type.

| Material Type | Coated | Uncoated | Total |
| --- | --- | --- | --- |
| CTB | 0 | 1 | 1 |
| E-leather | 1 | 2 | 3 |
| Gauze sponge | 0 | 1 | 1 |
| Seat belt strap | 2 | 2 | 4 |
| Tray table | 0 | 2 | 2 |
| **Total** | **3** | **8** | **11** |

The mean (x□) bacterial CFU/cm^2^ across all coupons was 354 CFU/cm^2^ with a standard deviation (σ) of 566.17 CFU/cm^2^. Coated coupons had a lower mean bacterial CFU/cm^2^ (x□ = 85.5, σ = 159.4) than that of uncoated coupons (x□ = 622.5, σ = 700.8). This difference was found to be statistically significant (Fig. 3), where the coated coupons had 1.387 log_10_, or 95.9%, reduction in bacterial CFU/cm^2^ when compared with the bacterial CFU/cm^2^ on the uncoated coupons (β = -1.387, Confidence Interval (CI, lower-upper) = -2.519 - -0.256, degrees of freedom (*df*) = 18, *p-value* = 0.019). Among coupon material types, there was no statistically significant difference in log_10_ bacterial CFU/cm^2^ (ANOVA, F = 1.61, df = 4, *p-value* = 0.223) (S1 Fig. 1).

There were eight bacterial genera found to make up five percent or more of the total abundance each by CFU/cm^2^(Table 3). Three genera dominated the bacterial community abundance, *Neisseria* sp.*, Streptococcus* sp. and unidentifiable bacterium. Among fungal isolates, only *Penicillium* sp. was present (which was also found on the Galley – 1 placard). While not present on the Galley – 3 placard, *Rhodotorula* sp. was found to be present on the exercise area placard. All fungi were found in low abundance, at approximately 15 CFU/cm^2^.

**Fig. 3.**
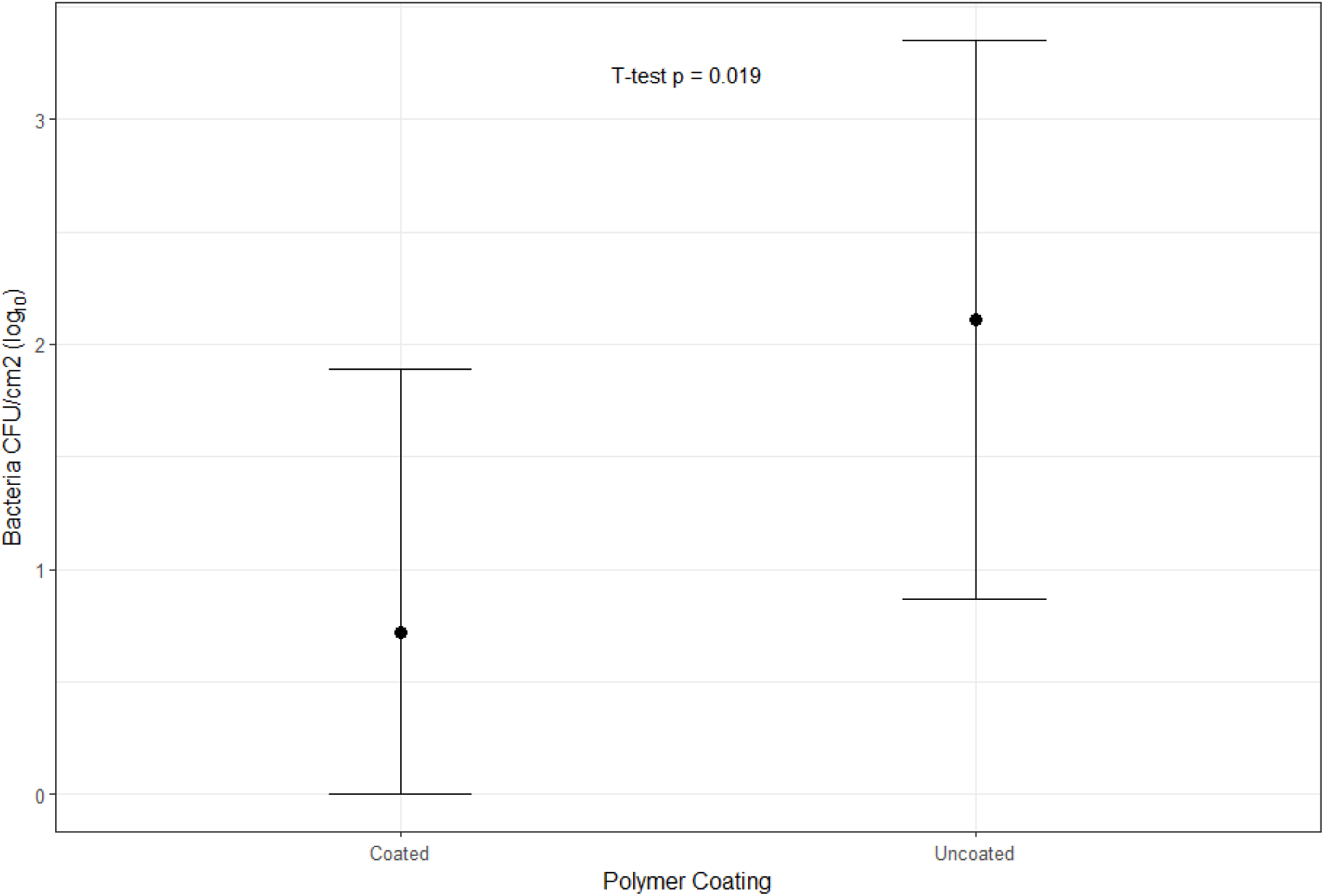
Mean log_10_ bacterial CFU/cm^2^ (black point) and 95% confidence intervals (bars) and statistical T-test *p-value* for coated and uncoated materials.

**Table 3.** Percentage (%) abundance of total identified CFUs and genetic material above 5% from bacterial cultures and genetic qPCR, respectively.

| Unique Species | Culture | Genetic qPCR |
| --- | --- | --- |
| *Gemella haemolysans* | 8.1 | - |
| *Streptococcus* sp.<sup>#</sup> | 5.0 | - |
| *Neisseria* sp. | 28.4 | - |
| *Pseudomonas* sp. | 6.9 | - |
| *Rothia* sp. | 6.2 | - |
| *Staphylococcus* sp. | 7.2 | 18.4 |
| *Streptococcus* sp. | 17.4 | 5.8 |
| *Corynebacterium* sp. | - | 13.5 |
| *Cutibacterium* sp. | - | 30.2 |
| Other/Unidentified bacterium | 20.8 | 32.1 |

\* The "Culture" column entries are percent abundance calculated from identified CFU counts recovered via culture (CFU/cm<sup>2</sup> normalized across positive coupons) from the Galley – 3 placard. The "Genetic qPCR" column entries are the percent contribution of taxa to the total 16S gene copy signal as measured by the assay from the hygiene placard. <sup>#</sup> Are the organisms that were unable to be identified to genus level.

### Microbial Communities

The microbial species communities on the coated and uncoated coupon surfaces were found not to be statistically significant between one another (ANOSIM R = -0.04, *p-value* = 0.914), nor among the different material types (ANOSIM *R* = -0.02, *p-value* = 0.662). Supporting this, the variance in microbial species communities were further not statistically explained by coated and uncoated samples (PERMANOVA *R2* = 0.024, *F* = 0.675, *p-value* = 0.934), or by material type (PERMANOVA *R2* = 0.1153, *F* = 0.815, *p-value* = 0.944). Assessing the homogeneity of multivariate dispersions showed that the average distance to group centroid was very similar for coated versus uncoated coupons (mean distance: Coated = 0.340, Uncoated = 0.322) and that this difference was not significant (ANOVA *F* = 0.311, *p-value* = 0.581). Similarly, dispersion among material groups did not differ significantly (mean distances: CTB = 0.311, E-leather = 0.337, Gauze sponge = 0.291, Seat belt strap = 0.298, Tray table = 0.324; ANOVA *F* = 0.310, *p-value* = 0.868). These results suggest there are no detectable changes in community centroid location or within-group dispersion attributable to coating or material type from the hygiene placard.

Whilst the community of species didn’t differ, there were statistically significant differences in the amount of genetic material per species. On the uncoated coupons, using multilevel pattern analysis, there were two species that were found to be strongly and statistically significantly associated with the uncoated coupons; *Brevibacterium* sp. (MPA *stat* = 0.44, *p-value* = 0.004) and *Fusobacterium* sp. (*stat* = 0.37, *p-value* = 0.033).

#### Species Abundance

From the genomic analysis on the coupons of the hygiene placard, we found that for all materials, except for the gauze sponge and an a single coupon outlier on the E-leather, the uncoated coupons had on average a greater abundance of genetic material than that of the coated coupons (Fig. 4).

**Fig. 4.**
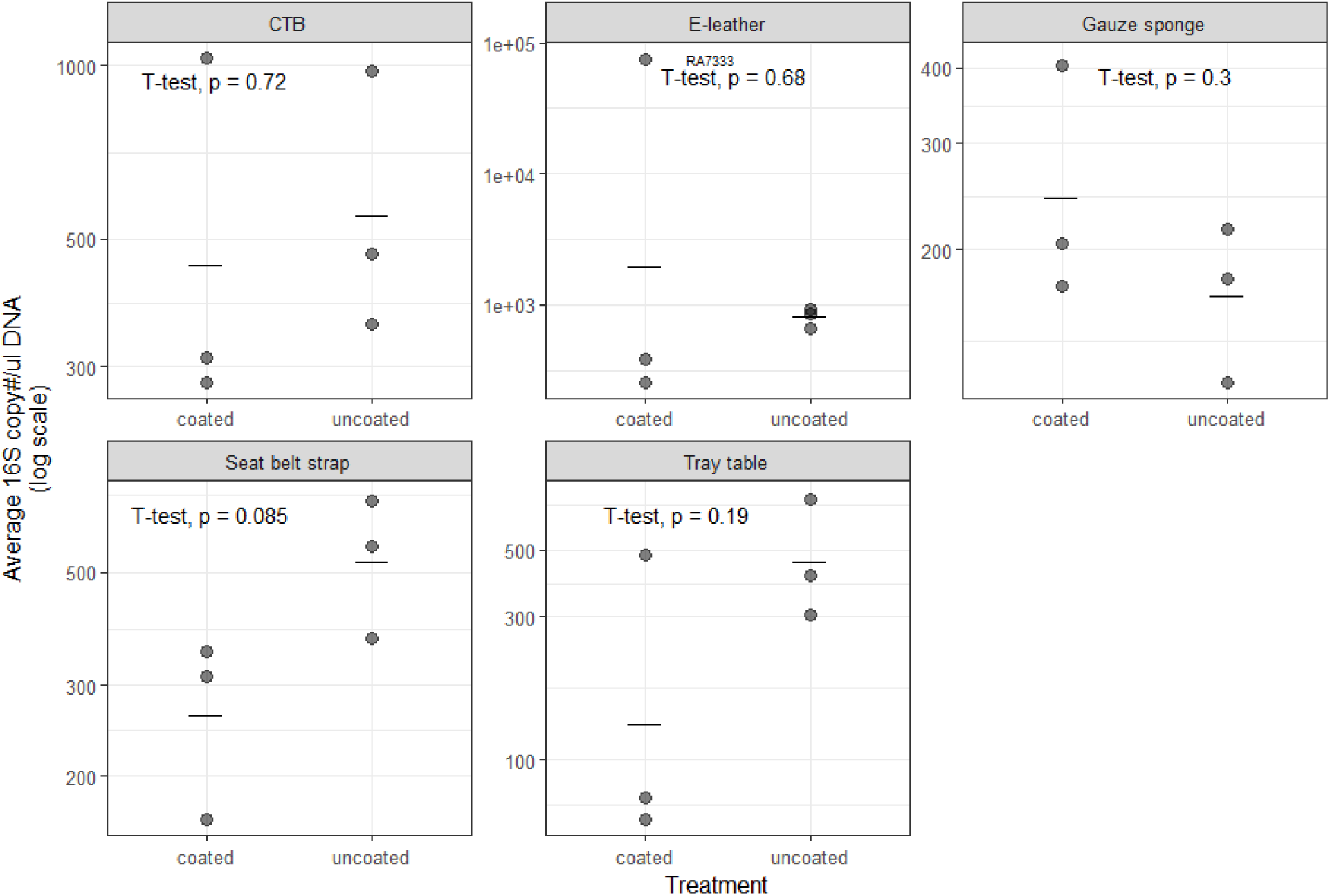
Observed abundance of bacterial 16S SSU rRNA gene copies by polymer coating and material type as measured by qPCR. The horizontal lines show the geometric mean of each group to mirror the results from the analysis with a log-transformed response variable and the *p-value* from a t-test for significance between these means.

The polymer coating and material type both had a statistically significant difference in microbial abundance (*F =* 6.4, *df* = 1,19, *p-value* = 0.020) and (*F =* 3.4, *df* = 4,19, *p-value* = 0.030) respectively, however; no statistically significant association was seen between the coating and material types (*F =* 2.0, *df* = 4,19, *p-value* = 0.143). The difference in material type was primarily driven by the tray table material having the lowest median microbial abundance (Ratio median = 0.29, CI = 0.11-0.74, *t* = -2.7, *df* = 19, *p-value* = 0.013) (S1 Table 3).

Genetic cycle threshold (Ct) values were generally high overall with a low number of gene copies. A summary of the observed Ct results by polymer coating and material type for the neat dilution is shown in supporting information 1 Table 4. Species richness was, on average, seen to be greater in uncoated samples across material types, except for E-leather and gauze sponge (Fig. 5), although this was not statistically significant. Moreover, material type showed a significant difference in number of species even after controlling for the outlier sample in E-leather (Table 4).

**Fig. 5.**
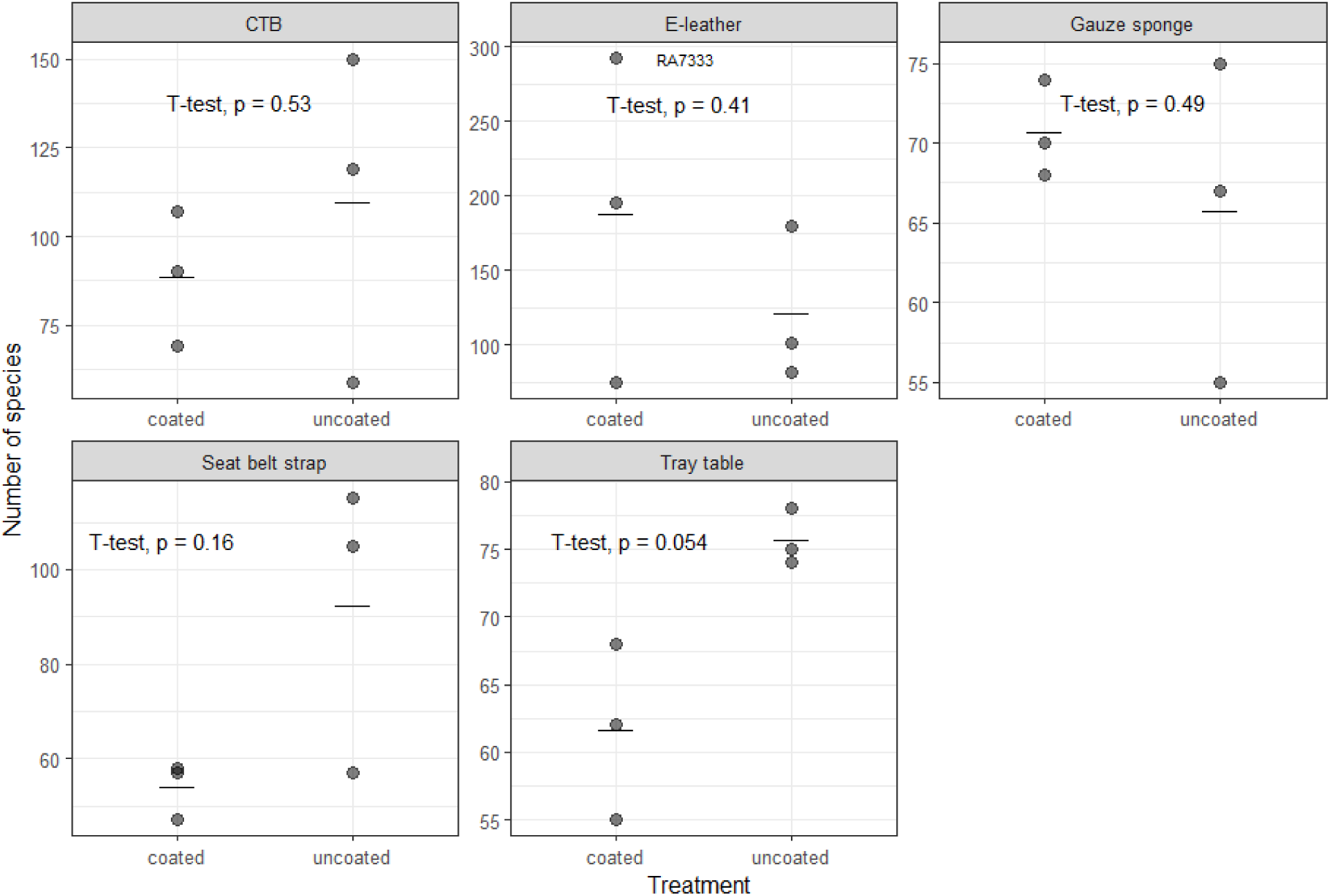
Observed richness values by treatment and material type plotted as points. The horizontal lines show the mean of each group and the *p-value* from a t-test for significance between these means. Each pane was allowed to have a different y axis scale. The outlier sample on the e-leather material is labeled with the sample ID.

**Table 4.** The type of material and its effect on species richness with and without the outlier sample from the E-leather material.

| Effect | $\chi^2$ | df | p-value |
| --- | --- | --- | --- |
| Without outlier | 27.3 | 4 | < 0.0001 |
| With outlier | 43.7 | 4 | < 0.0001 |

### Polymer Durability

The polymer coating on the coupons from the Galley – 2 placard were found to remain on the surface of all material types regardless of whether they were touched throughout the experimental period or not (t-test *t* = 0.909, *df* = 27.841, *p-value* = 0.371), with the touched surfaces having a greater mean gray value than that of the controls on all material types (Fig. 6), however this difference was not statistically significant. Moreover, when taking bi-weekly measurements of the ground samples, there was no observable decline in the amount of polymer coating present on the surface of the touched surfaces compared with the control surfaces (Fig. 7).

**Fig. 6.**
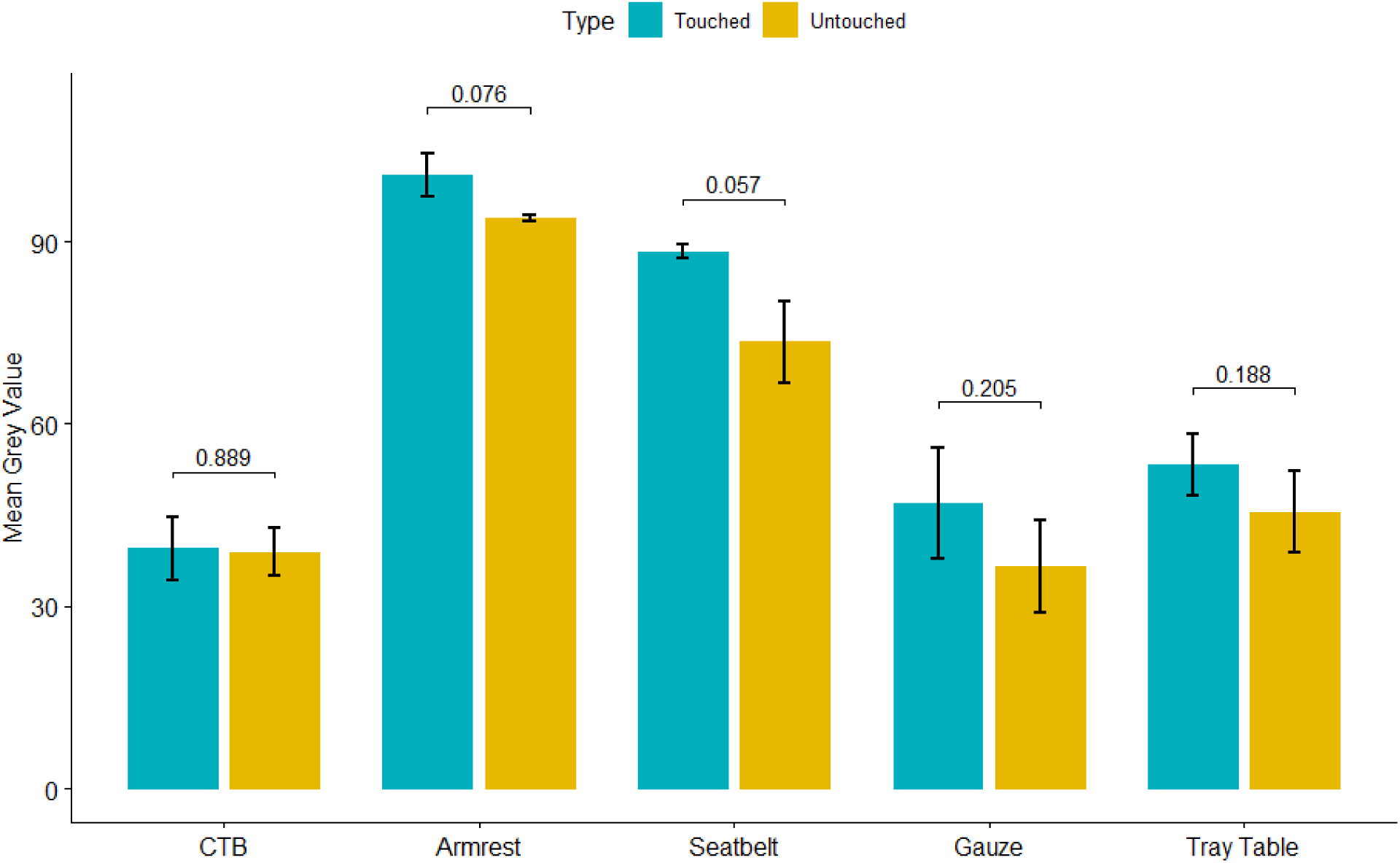
Mean and standard deviation of the mean grey value recorded form the fluorescence showing the remaining polymer from ISS surfaces by material type and exposure to touching. Within group comparisons of the mean grew value is given above the bars using a t-test, showing the *p-value* statistical significance.

**Fig. 7.**
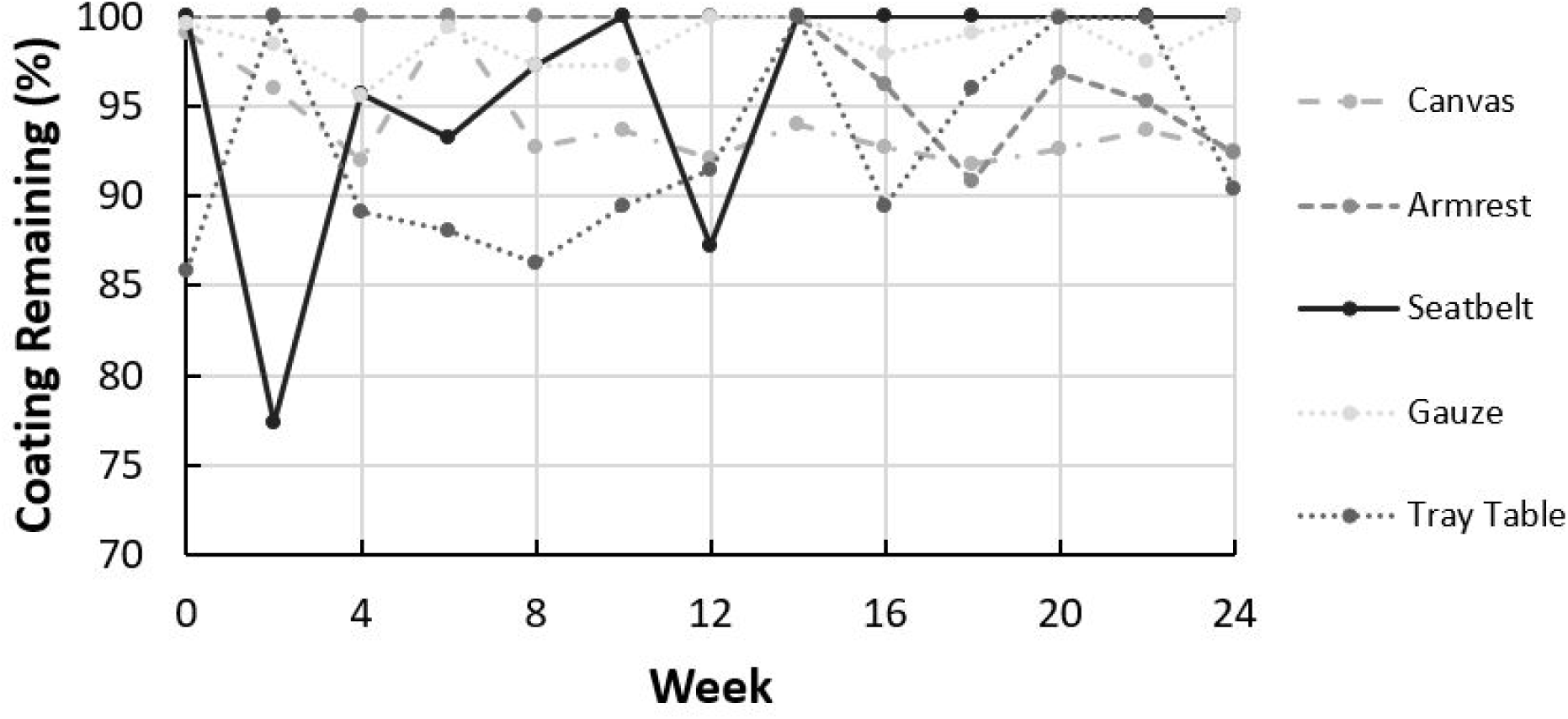
Longevity of polymer coating between touched and control surfaces throughout time on ground samples.

## Discussion

Microorganisms are ubiquitous, being present in nearly all natural and built environments, including those in space. Microbes pose a challenge to space flight, as pathogenic microbes can impact crew health and environmental microbes can impact upon the spacecraft when they degrade vehicle materials or interfere with systems function. We explored microbial presence on different surfaces aboard the ISS from different material types and tested the efficacy of a novel antimicrobial polymer. Here we examine a new previously described antimicrobial polymer^15,18^ and its efficacy and durability in spacecraft in zero gravity aboard the ISS over a six-month period. The findings from our study provide valuable insights into the microbial community aboard the ISS and the efficacy of this novel antimicrobial polymer coating in mitigating microbial growth. We found that microbial growth across all sampling locations was limited, and where there were microbes present, our polymer coating was able to reduce microbial load and growth.

Among the placards that underwent microbial culturing, there was only one placard, Galley - 3 placard, that exhibited sufficient microbial loads across the coupons to conduct and determine a statistically significant reduction in microbial burden on the coated samples, achieving a 1.387 log_10_ reduction in bacterial CFU/cm². This represents an approximate 95.9% reduction in bacterial presence on surfaces coated with the polymer. This supports previous testing of the polymer against viruses, such as influenza, in Earth based observational studies where it was shown to significantly reduce viral loads[14]. The most growth observed on a single coupon was in the uncoated tray table material with 320 CFU/cm², which is, while higher, comparable to the 30 CFU/cm² reported in earlier studies[31]. On the Galley - 3 placard, the polymer coating was seen to reduced viable bacterial recovery, indicating that the coating can be effective under low microbial load, however, broader generalization to other locations requires additional testing.

We expected that the non-cultivation-based methods would detect a high abundance and diversity of microorganisms present on the coupons. However, the genetic analysis conducted on the hygiene placard showed similar minimal growth seen in microbiological cultures, confirming our initial finding of limited microbial buildup on the test coupon surface aboard the ISS.The cycle threshold (Ct) values from qPCR were consistently high, which could imply bacterial deposition but not survival for most areas of the ISS. The overall amount of genetic material differed by coupon material type. Notably, the seat belt material had statistically significantly greater genetic material abundance than compared to the tray table material. This difference may be attributed to the texture, absorption, and physical properties of the material where microbes may penetrate deeper into porous surfaces or have available metabolites to promote growth. However, we found no statistically significant difference in bacterial CFU/cm^2^ by material type. We originally expected the hygiene placard to show greater aerosolized particles due to exfoliation behaviors[32], yet this was not observed; this could be attributed to the higher air filtration and flow rates on the spacecraft, a paradigm analogous to aircraft[10].

We observed a large amount of species’ richness in our samples, including an outlier from one sample having both high levels of DNA and species richness. This suggests that the microbial community is not merely a result of a single CFU expanding but rather represents a diverse community of species. The polymer’s durability further showed minimal degradation of the coating over the six-month study period. However, if the polymer is applied in conjunction with cleaning, an adhesion promoter may be required to counteract its potential removal, though we note part of the goal of the polymer’s presence is to avoid cleaning. Due to the low bacterial presence and genetic read counts from the aerosol placards, we cannot make definitive comments on the correlation between microbial load and durability. Future studies could explore this relationship further, as while we feel it is unlikely, higher microbial loads could inundate the polymer and lower the polymer’s efficacy.

The lack of culturable organisms on the fomite placard raises questions about the potential for self-cleaning during surface touch events. One possibility is that the microbes deposited via touch were subsequently ingested, spread around the ISS environment, or potentially concentrated at specific areas where we did not have placards The positioning of the placards on the ISS may have influenced the microbial communities observed, with potential sources including foodborne and fecal bacteria, and microbes from the human microbiome. The bacteria identified were generally not pathogenic; however, they may still pose potential concerns for astronauts on long-term missions, where immunosuppression could exacerbate risks. Notably, one of the two uncultured bacteria significantly associated with the uncoated communicated were Gram-positive, with one being aerobic and the other an anaerobic species. The presence of *Fusobacterium* sp. is notable because this genus comprises obligate anaerobes that are more commonly associated with the human oral cavity and gastrointestinal tract than with indoor environments. Its presence more likely reflects crew-associated deposition or transport for airborne particulate matter than from sustained growth on cabin surfaces. However, it is an interesting finding given its detection was on one of the aerosols (untouched) placards, which supports the idea that its deposition was from airborne bound particulate matter. The presence of the *N. gonorrhoeae* genus, which includes pathogenic species such as *Neisseria* gonorrhoeae, is particularly noteworthy, as many species within this genus are primarily soil-dwelling and can be infectious. We also note the presence of microbial species associated with material degradation and biofilm formation, including *Penicillium*, which pose risks to equipment longevity[5].

Our findings align with previous studies that have reported *Staphylococcus* and *Streptococcus* onboard space craft surfaces[33]. The presence of *Penicillium*, which was tied for the highest abundance of yeast with *Rhodotorula* in our analysis, supports findings from other studies as leading contributors to the microbial community abundance[34]. While in Venkateswaran et al. (2014), more than 90% of all genetic sequences were assigned to four bacterial genera (*Corynebacterium, Propionibacterium, Staphylococcus,* and *Streptococcus*), in our study, these genera comprised only 34.7% of the data (10.5%, 0%, 18.4%, and 5.8%, respectively)[35]. This difference highlights the unique microbial ecology of the ISS, its potential to change through time in a closed-loop system, and for microbial adaptation in space.

The source of microbes aboard the ISS remains a topic of interest for crew health, environmental systems, food production, and vehicle safety. Previous studies have suggested that the microbial communities on the ISS are more similar to those found in human homes rather than to those on the human body[36]. Our findings support this, as we identified environmental bacteria such as *Neisseria* and *Staphylococcus*, which are likely to originate from clothing fibers, skin, food, and other sources[32]. In the absence of gravitational settling of microbes onto surfaces, the ISS may allow larger particles to persist in the internal atmosphere in areas where air flow doesn’t readily move particles to the air filtration system, creating unique exposure environments for astronauts. The lack of microbial abundance found here may have future impacts on research and operational protocols aboard the ISS. For instance, the lack of microbes could reflect the cleaning efficacy by the crew and ventilation system, and the potential of overcleaning, where there may be a need to balance maintaining a sterile environment and the practicalities of cleaning protocols, reducing the valuable time crew required for cleaning. Despite the low bacterial counts found, the closed environment of the ISS may present niche microbial challenges where hotspots for microbial accumulation may exist via touch or aerosol deposition and were simply not incorporated in this study. As an example, the positioning of placards, such as Galley – 3, which was located above other galley placards, could verify airflow dependent microbial deposition. Future air sampling and modeling simulations could aid in predicting the optimal placement for antimicrobial interventions, particularly in high-traffic areas.

The polymer has undergone several methods of testing regarding determining its toxicity and safety. This has included being classified as a polymer of low concern by the Australian Industrial Chemicals Introduction Scheme and via testing being conducted by Eurofins using the US Environmental Protection Agency’s Test Guidelines for Pesticides and Toxic Substances [unpublished, in preparation for publication]. However, one limitation of our study is the potential selection for highly adapted microbes aboard the ISS, such as spore formers, which our polymer may not effectively target[5]. Additionally, the low biomass challenge we faced, coupled with the different methods used in our culture laboratory, may have influenced our results[37]. The tightly controlled cabin humidity aboard the ISS may have contributed to the low bacterial presence observed on the surface coupons. With low moisture levels, there is limited moisture deposition on the coupons, which further inhibits active microbial growth and proliferation. The logistics of space-related payloads impacted our approach and sampling procedures, including the number of samples and replicates taken, as well as the timing of sample collection[5]. Previous efforts to detect microbes in space have shown that while microbiological cultures assist in revealing post-flight microbial viability, inherent issues with microbial die-off during sample return have caused genomic analysis to have a mainstay-place in analysis and assessment of microbial levels. Here we applied microbiological culturing and genomic analysis to capture both viable and nonviable organisms and have findings representative of what microbes circulate in the ISS.

A limitation of the genomic dataset used here is that it was derived from a single coupon set per polymer coating and material combination rather than multiple biological replicates, which limits our ability to quantify within-group variability and evaluate the influence of outliers more robustly. In addition, low-biomass surface samples are particularly susceptible to contamination during collection, extraction, and amplification. To address this, we processed blank extraction and packaging controls in parallel and screened taxa/features against these negative controls during data interpretation. Although this approach supports confidence that the dominant signals observed in the hygiene placard dataset were not solely attributable to contamination, the low biomass of the samples means that the genomic results should still be interpreted with caution. Future flight studies with replicated coupon sets and matched sequencing blanks will be important to confirm these findings and improve estimation of coating effects on the ISS microbiome.

Our study highlights the low microbial counts observed from natural environmental deposition and from the touching interaction from the astronaut crew, with deposited bacteria not actively replicating at high levels. We show an approximate 95.9% reduction in bacterial presence on surfaces with the use of the polymer coating. The durability and presence of the polymer persisted the entire duration of the six-month experimental period, indicating its potential for long-term use in space environments. The mitigation of microbial contamination in space craft requires a “Swiss cheese” multi-layered approach, combining multiple technologies and strategies. The findings of the polymers efficacy and durability longevity on surfaces may support its use in these multi-layered risk reduction systems, providing long lasting reduction in microbial loads. Future work should focus on evaluating hotspots based on airflow, crew contact, and swabbing, followed by assessing this polymer coating as an intervention in these targeted areas. Additionally, understanding the role of touch in the ISS context may provide insights into microbial removal and cleaning efficacy. This may further our understanding of how we may optimize cleaning regimes aboard crewed space vehicles. Overall, our findings contribute to the ongoing efforts to ensure astronaut health and safety in the unique environment of space.

## Acknowledgments

The authors wish to thank Scott Copeland, Mohammad Abbas, Melissa Boyer, and Carlos Sahagun who were the liaison with the International Space Station Program for all technical communication and assistance with the deployment of the experiment. We would further like to thank and acknowledge all the International Space Station Program ground and space crew and employees who carried out the experimental setup, training, and conducting the experiment.

## Author Contributions

**Conceptualization:** Jason Armstrong, Iain Koolhof, David Corporal, Mark Wilson, Michael Monteiro

**Data Curation:** Iain Koolhof, David Corporal, Olivia Jessop, Mark Wilson, Darren Dunlap

**Formal Analysis:** Iain Koolhof, Olivia Jessop, Ariel Muldoon, Mark Wilson, Darren Dunlap

**Investigation:** David Corporal, Mark Wilson, Darren Dunlap

**Methodology:** Jason Armstrong, Iain Koolhof, David Corporal, Olivia Jessop, Mark Wilson, Darren Dunlap, Michael Monteiro

**Project Administration:** Jason Armstrong, Iain Koolhof, David Corporal, Mark Wilson, Michael Monteiro

**Resources:** Jason Armstrong, David Corporal, Mark Wilson, Michael Monteiro

**Software:** Iain Koolhof, Ariel Muldoon

**Supervision:** Jason Armstrong, Reuben Strydom, Iain Koolhof, David Corporal, Michael Monteiro

**Visualization:** Iain Koolhof, Ariel Muldoon

**Writing – Original Draft:** Iain Koolhof, David Corporal, Olivia Jessop, Mark Wilson, Ariel Muldoon, Darren Dunlap, Sung-Po Chen, Nicholle Wallwork

**Writing – Review & Editing:** Jason Armstrong, Iain Koolhof, David Corporal, Sung-Po Chen, Nicholle Wallwork, Reuben Strydom, Mark Wilson, Darren Dunlap, Olivia Jessop, Ariel Muldoon, Michael Monteiro

## References

1. Armstrong JW, Gerren RA, Chapes SK. The effect of space and parabolic flight on macrophage hematopoiesis and function. Exp Cell Res. 1995;216: 160–168.

2. Bijlani S, Stephens E, Singh NK, Venkateswaran K, Wang CCC. Advances in space microbiology. iScience. 2021;24: 102395. doi:10.1016/j.isci.2021.102395

3. Pierson D, Bruce R, Ott CM, Castro V, Mehta S. Microbiological Lessons Learned From the Space Shuttle. 41st International Conference on Environmental Systems. Portland, Oregon: American Institute of Aeronautics and Astronautics; 2011. doi:10.2514/6.2011-5266

4. Paton S, Moore G, Campagnolo L, Pottage T. Antimicrobial surfaces for use on inhabited space craft: A review. Life Sci Space Res. 2020;26: 125–131. doi:10.1016/j.lssr.2020.05.004

5. Mora M, Wink L, Kögler I, Mahnert A, Rettberg P, Schwendner P, et al. Space Station conditions are selective but do not alter microbial characteristics relevant to human health. Nat Commun. 2019;10: 3990. doi:10.1038/s41467-019-11682-z

6. Szydlowski LM, Bulbul AA, Simpson AC, Kaya DE, Singh NK, Sezerman UO, et al. Adaptation to space conditions of novel bacterial species isolated from the International Space Station revealed by functional gene annotations and comparative genome analysis. Microbiome. 2024;12: 190. doi:10.1186/s40168-024-01916-8

7. Simpson AC, Sengupta P, Zhang F, Hameed A, Parker CW, Singh NK, et al. Phylogenomics, phenotypic, and functional traits of five novel (Earth-derived) bacterial species isolated from the International Space Station and their prevalence in metagenomes. Sci Rep. 2023;13: 19207. doi:10.1038/s41598-023-44172-w

8. Wang M, Duday D, Scolan E, Perbal S, Prato M, Lasseur C, et al. Antimicrobial Surfaces for Applications on Confined Inhabited Space Stations. Adv Mater Interfaces. 2021;8: 2100118. doi:10.1002/admi.202100118

9. Trent S, Davis A, Wu T, Menard D, Cummins J, Santarpia J, et al. Inhaled Mass and Particle Removal Dynamics in Commercial Buildings And Aircraft Cabins. Ashrae. 2022;64.

10. Pang JK, Jones SP, Waite LL, Olson NA, Armstrong JW, Atmur RJ, et al. Probability and estimated risk of SARS-CoV-2 transmission in the air travel system. Travel Med Infect Dis. 2021;43: 102133. doi:10.1016/j.tmaid.2021.102133

11. Bolyen E, Rideout JR, Dillon MR, Bokulich NA, Abnet CC, Al-Ghalith GA, et al. Author Correction: Reproducible, interactive, scalable and extensible microbiome data science using QIIME 2. Nat Biotechnol. 2019;37: 1091–1091. doi:10.1038/s41587-019-0252-6

12. Page K, Wilson M, Parkin IP. Antimicrobial surfaces and their potential in reducing the role of the inanimate environment in the incidence of hospital-acquired infections. J Mater Chem. 2009;19: 3819–3831.

13. Vélez Justiniano Y-A, Goeres DM, Sandvik EL, Kjellerup BV, Sysoeva TA, Harris JS, et al. Mitigation and use of biofilms in space for the benefit of human space exploration. Biofilm. 2023;5: 100102. doi:10.1016/j.bioflm.2022.100102

14. Bobrin VA, Chen S-P, Grandes Reyes CF, Sun B, Ng CK, Kim Y, et al. Water-Borne Nanocoating for Rapid Inactivation of SARS-CoV-2 and Other Viruses. ACS Nano. 2021;15: 14915–14927.

15. Zea L, McLean RJC, Rook TA, Angle G, Carter DL, Delegard A, et al. Potential biofilm control strategies for extended spaceflight missions. Biofilm. 2020;2: 100026. doi:10.1016/j.bioflm.2020.100026

16. Lemelle L, Campagnolo L, Mottin E, Le Tourneau D, Garre E, Marcoux P, et al. Towards a passive limitation of particle surface contamination in the Columbus module (ISS) during the MATISS experiment of the Proxima Mission. Npj Microgravity. 2020;6: 29. doi:10.1038/s41526-020-00120-w

17. Bobrin VA, Chen S-PR, Grandes Reyes CF, Smith T, Purcell DFJ, Armstrong J, et al. Surface Inactivation of Highly Mutated SARS-CoV-2 Variants of Concern: Alpha, Delta, and Omicron. Biomacromolecules. 2022;23: 3960–3967. doi:10.1021/acs.biomac.2c00801

18. Turner J. Boeing Environment Responding Antimicrobial Coatings. In: Space Station Research Explorer [Internet]. [cited 9 Dec 2024]. Available: https://www.nasa.gov/mission/station/research-explorer/investigation/?#

19. Turner J. International Space Station Boeing Antimicrobial Coating. In: Space Station Research Explorer [Internet]. [cited 9 Dec 2024]. Available: https://www.nasa.gov/mission/station/research-explorer/investigation/?#id=8946

20. Rice EW, Bridgewater L, American Public Health Association. Standard methods for the examination of water and wastewater. American public health association Washington, DC; 2012.

21. Yamaguchi N, Roberts M, Castro S, Oubre C, Makimura K, Leys N, et al. Microbial Monitoring of Crewed Habitats in Space—Current Status and Future Perspectives. Microbes Environ. 2014;29: 250–260. doi:10.1264/jsme2.ME14031

22. Ludwig W. ARB: a software environment for sequence data. Nucleic Acids Res. 2004;32: 1363–1371. doi:10.1093/nar/gkh293

23. Minchin PR. An evaluation of the relative robustness of techniques for ecological ordination. Springer; 1987. pp. 89–107.

24. R Core Team. R: A Language and Environment for Statistical Computing. R Foundation for Statistical Computing; 2021. Available: https://www.R-project.org/

25. Venables WN, Ripley BD. Modern Applied Statistics with S. Fourth. New York: Springer; 2002. Available: https://www.stats.ox.ac.uk/pub/MASS4/

26. Oksanen J, Kindt R, Legendre P, O’Hara B, Simpson GL, Solymos P, et al. vegan: Community Ecology Package. 2008. Available: http://cran.r-project.org/, http://vegan.r-forge.r-project.org/

27. Wickham H. ggplot2: Elegant Graphics for Data Analysis. Springer-Verlag New York; 2016. Available: https://ggplot2.tidyverse.org

28. Hartig F. DHARMa: Residual Diagnostics for Hierarchical (Multi-Level / Mixed) Regression Models. 2024. Available: https://CRAN.R-project.org/package=DHARMa

29. Cáceres MD, Legendre P. Associations between species and groups of sites: indices and statistical inference. Ecology. 2009;90: 3566–3574. doi:10.1890/08-1823.1

30. RStudio Team. RStudio: Integrated Development Environment for R. Boston, MA: RStudio, PBC.; 2020. Available: http://www.rstudio.com/

31. Urbaniak C, Morrison MD, Thissen JB, Karouia F, Smith DJ, Mehta S, et al. Microbial Tracking-2, a metagenomics analysis of bacteria and fungi onboard the International Space Station. Microbiome. 2022;10: 100. doi:10.1186/s40168-022-01293-0

32. Reuland CL, Brown CA. Evaluation of the Accumulation of Foreign Object Debris in the International Space Station Ventilation Systems and Resulting Impacts to Systems.

33. Taylor PW. Impact of space flight on bacterial virulence and antibiotic susceptibility. Infect Drug Resist. 2015; 249–262.

34. Benardini III JN, La Duc MT, Ballou D, Koukol R. Implementing planetary protection on the atlas v fairing and ground systems used to launch the Mars Science Laboratory. Astrobiology. 2014;14: 33–41.

35. Venkateswaran K, Vaishampayan P, Cisneros J, Pierson DL, Rogers SO, Perry J. International Space Station environmental microbiome — microbial inventories of ISS filter debris. Appl Microbiol Biotechnol. 2014;98: 6453–6466. doi:10.1007/s00253-014-5650-6

36. Lang JM, Coil DA, Neches RY, Brown WE, Cavalier D, Severance M, et al. A microbial survey of the International Space Station (ISS). PeerJ. 2017;5: e4029. doi:10.7717/peerj.4029

37. Stahl-Rommel S, Jain M, Nguyen HN, Arnold RR, Aunon-Chancellor SM, Sharp GM, et al. Real-Time Culture-Independent Microbial Profiling Onboard the International Space Station Using Nanopore Sequencing. Genes. 2021;12: 106. doi:10.3390/genes12010106

