## supporting information 1 for "A next generation space antimicrobial: assessing microbial activity and reduction across the International Space Station"

***Tables***

Table 1. Primers used for laboratory sequencing identification^1,2^

| Purpose | Primer |
| --- | --- |
| Bacterial | 27F: AGAGTTTGATCCTGGCTCAG |
|  | 1492R: GGTTACCTTGTTACGACTT |
| Fungal | ITS1: TCCGTWGGTGAACCWGCGG |
|  | NL4: GGTCCGTGTTTCAAGACGGG |

Table 2. Number of coupons that cultured bacterial growth per placard

| **Placard** | **Total instance of growth from unique coupons** | **Instances of growth including replicates** |
| --- | --- | --- |
| Exercise | 2 | 2 |
| Galley 1 | 3 | 4 |
| Galley 3 | 11 | 19 |
| Toilet | 2 | 3 |

Table 3: Estimated ratios of medians, with 95% confidence intervals, from the fitted model. Estimated differences are median coated treatment abundance divided by median uncoated treatment abundance. The overall treatment effect is labeled as "Overall".

| Material type | Ratio of medians (coated/uncoated) | 95% CI | *t* | df | *p-value* |
| --- | --- | --- | --- | --- | --- |
| Overall | 0.58 | 0.38, 0.90 | -2.6 | 19 | 0.0174 |
| CTB | 0.82 | 0.32, 2.10 | -0.4 | 19 | 0.6645 |
| E-leather | 0.39 | 0.14, 1.13 | -1.9 | 19 | 0.0790 |
| Gauze sponge | 1.45 | 0.57, 3.73 | 0.8 | 19 | 0.4173 |
| Seat belt strap | 0.50 | 0.20, 1.28 | -1.5 | 19 | 0.1406 |
| Tray table | 0.29 | 0.11, 0.75 | -2.7 | 19 | 0.0128 |

Table 4: A summary of observed Ct values for the neat dilution for each material type and treatment. This includes the total number of replicates with non-missing Ct (maximum 9) plus the mean and range.

| Material type | Treatment | Total replicates with Ct | Mean Ct | Range Ct |
| --- | --- | --- | --- | --- |
| CTB | coated | 4 | 36.0 | 34.3 - 36.9 |
|  | uncoated | 5 | 35.3 | 34.3 - 36.2 |
| E-leather | coated | 5 | 34.5 | 26.4 - 37.1 |
|  | uncoated | 6 | 34.7 | 34.3 - 35.3 |
| Gauze sponge | coated | 5 | 36.8 | 35.8 - 37.5 |
|  | uncoated | 5 | 37.6 | 37.1 - 38.7 |
| Seat belt strap | coated | 4 | 36.7 | 36.2 - 37.6 |
|  | uncoated | 6 | 35.5 | 34.9 - 36.4 |
| Tray table | coated | 5 | 37.9 | 35.2 - 39.4 |
|  | uncoated | 7 | 35.8 | 34.7 - 36.7 |

***Figures***

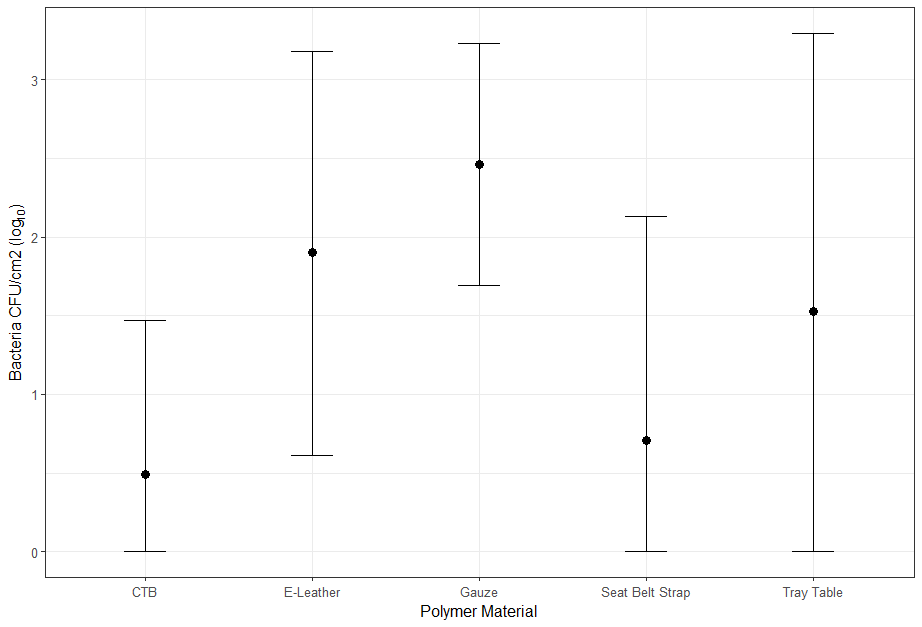

**Fig. 1.** Mean log_10_ bacterial CFU/cm^2^ (black point) and 95% confidence intervals (bars) per material type.

***Methods***

### Polymer Mixture Preparation and Sterilization methods

The polymer latex (10 wt.%, 40 g of latex, 4 g of polymer and 3.63 $\times$ 10^-3^ mol of DMAEMA units) was diluted with 28 mL of water, then propargyl bromide (80 wt.% in toluene, 27 mg, 1.81 $\times$ 10^-4^ mol) in 16 mL of ethanol was added and the quaternization reaction proceeded at 25 °C for 3 h. To the quaternized polymer latex, 3-azido-7-hydroxycoumarin (36.9 mg, 1.81 $\times$ 10^-4^ mol) in 8.7 mL of ethanol was added, and the resulting dispersion was degassed with argon for 60 min. Ascorbic acid (0.1917 g, 1.09 $\times$ 10^-3^ mol) was dissolved in 5 mL of water, degassed with argon for 30 min, and injected into the reaction mixture. Copper sulfate pentahydrate (0.1737 g, 1.09 $\times$ 10^-3^ mol) was dissolved in 5 mL of water, degassed with argon for 30 min, and injected into the reaction mixture. The CuAAC reaction was stirred under argon overnight, then purified by dialysis (10k MWCO) against water (6 $\times$ 2 L) over 72 h. The resulting nanoworm dispersion filtered through cotton plug to remove larger impurities that remained after dialysis.

The hardware involved in the preparation of the samples was sterilized using either autoclave, UV light, or 70 % ethanol, depending on the tolerance of the material to the sterilization technique. The TOMY SX-700 autoclave operating under sterilize/normal function at 121 °C for 20 min was used to sterilize the placard containers, placards, container lids, seat belt strap coupons, cargo bag coupons, cotton sponge coupons, and tray table coupons. The TOMY SX-700 autoclave operating under sterilize/liquid function at 121 °C for 30 min was used to sterilize a 2 L schott bottle containing 1.6 L of Milli-Q water. The Safemate Cyto 1.2 laminar flow cabinet equipped with a 30W UV-C tubular T8 lamp at 254 nm was used to sterilize the E-leather coupons, Velcro loops, Velcro hooks, ziplock bags, and stickers for 10 min. The remainder of the hardware and items were all sterilized by spraying with 70 % ethanol solution.

**References**

1.	Lane D i. Nucleic acid techniques in bacterial systematics. M. Goodfellow and A. G. O'Donnell (ed.), Handb New Bact Syst. 1991;115-147.

2.	Stackebrandt E., and W Liesack. Nucleic acids and classification. Handb New Bact Syst. 1993;151–189.
