## supporting information 2 for "A next generation space antimicrobial: assessing microbial activity and reduction across the International Space Station"

***Genetic Analysis***

### DNA extraction

The tip of each swab was snapped off into a 2 mL screw cap tube with O-ring (Sarstedt, cat #72.694.006) filled with 800 µL of Solution CD1 (supplied) and 300 mg 0.1 dia. glass beads (BioSpec Products, cat #11079101). DNA was lysed with a preliminary step of bead beating using the Powerlyser 24 homogenizer (Mo-Bio, #13155) for 5 min at 2000RPM. The tubes were heated at 65 ^0^C for 10 min, shaking at 1000 RPM, on the Eppendorf ThermoMixer (Eppendorf, cat #5382000031) and centrifuged for 1 min at 10 000 x g. The resulting lysate was transferred to a 1.5 mL Eppendorf tube (supplied). Samples were frozen and thawed to further lyse the cells. 200 µL of CD2 was added to the thawed lysate and extraction was as per Qiagen DNeasy Powersoil Pro Kit, (cat # 47016). Final elution volume was 50 µL.

### Quantitative PCR

From the eluted 50 µL DNA, qPCR was performed in triplicate at 3 dilutions (neat, 1:5, 1:10) using 6 μL of 2x QuantiNova SYBR Green PCR Master Mix (Qiagen, Germany), 4 µL template DNA and 1μL of primer mix (0.4 μM). The 16S primer set 1406F/1525R was designed to amplify bacterial and archaeal 16S rRNA genes (Table I). The PCR was run on the ViiA 7 Real Time PCR platform (Applied Biosystems, USA) using a cycle of 2 min at 95 °C and 40 cycles of [5 s at 95 °C and 20 s at 60 °C]. A melt curve was produced by running a cycle of 2 min at 95 °C and a last cycle of 15 s at 60 °C. The cycle threshold (Ct) values were recorded and analyzed using QuantStudio RealTime PCR software.

**Table 1**: qPCR primers

| Forward | Reverse |
| --- | --- |
| GYACWCACCGCCCGT | AAGGAGGTGWTCCARCC |

### Amplicon sequencing

Separately, the 16S rRNA gene encompassing the V3 and V4 regions was targeted using the 341F and

806R primers modified to contain Illumina-specific adapter sequence (Table II). The prokaryote primer pair Bac_SSU_341F-806R amplifies the small subunit (SSU) ribosomal RNA of bacteria (16S), specifically the V3 and V4 regions. In *E coli*, it amplifies the 341-806 region of the 16S gene.

Preparation of the 16S library was performed per the manufacturer guidelines (Illumina #15044223 Rev.B). In the 1st stage, PCR products of ~465 bp were amplified according to the specified workflow with an alteration in polymerase used to substitute NEBNext® Ultra™ II Q5® Mastermix (New England Biolabs #M0544) in standard PCR conditions. Resulting PCR amplicons were purified using Agencourt AMPure XP beads (Beckman Coulter, USA) at a ratio of 0.8x, so collecting fragments 500 bp, to avoid compromising the Q30 performance of the run, and possibly run failure.

Purified DNA was indexed with unique 8 bp barcodes using the Illumina Nextera XT 384 sample Index Kit A-D (Illumina FC-131-1002) in standard PCR conditions with NEBNext® Ultra™ II Q5® Mastermix (New England Biotech, USA). Indexed amplicons were pooled together in equimolar concentrations and sequenced on MiSeq Sequencing System (Illumina, USA) using paired end sequencing with V3 300 bp chemistry in the Australian Centre for Ecogenomics according to manufacturer’s protocol.

**Table 2**: Adapted 16S primers for DNA sequencing.

| Forward | Reverse |
| --- | --- |
| TCGTCGGCAGCGTCAGATGTGTATAAGAGACAGCCTACGGGNGGCWGCAG | GTCTCGTGGGCTCGGAGATGTGTATAAGAGACAGGACTACHVGGGTWTCTAATCC |

### Amplicon read processing and analysis pipeline

Passing quality control of resulting sequence is determined as 10,000 raw reads per sample prior to data processing and passing metrics in line with Illumina supplied reagent metrics of overall Q30 for 600 bp reads of >70%.

Primer sequences were removed from forward de-multiplexed reads using cutadapt (version 2.4)^23^, with reads not containing primers discarded (--discard-untrimmed). Poor quality reads were identified and removed with trimmomatic (version 0.39)^24^ using a sliding window of 4 bases with an average quality of 15 (SLIDINGWINDOW:4:15). Reads were cropped to 250 bp (CROP:250), with any less than 250 bp in length discarded (MINLEN:250). Only the forward reads we incorporated.

Quality controlled forward reads were processed using QIIME2 (ver. 2019.10.0)^12^ for feature selection, abundance calculations and taxonomy assignment. Reads were de-noised (filtered, dereplicated and chimeras identified and removed) using DADA2 (--p-trunc-len = 0)^25^ and relative frequencies of each resulting feature calculated. The taxonomy for each feature was assigned by BLASTing its sequence against the combined non-redundant 16S and 18S SILVA database (release 138, clustered at 99% identity)^26^ using the classify-consensus-blast function with default parameters.
